# The Tudor protein Veneno assembles the ping-pong amplification complex that 1 produces viral piRNAs in *Aedes* mosquitoes

**DOI:** 10.1101/242305

**Authors:** Joep Joosten, Pascal Miesen, Bas Pennings, Pascal W.T.C. Jansen, Martijn A. Huynen, Michiel Vermeulen, Ronald P. Van Rij

## Abstract

TUDOR-domain containing proteins facilitate PIWI interacting (pi)RNA biogenesis in *Drosophila melanogaster* and other model organisms. In *Aedes aegypti* mosquitoes, a somatically active piRNA pathway generates piRNAs from viral RNA during acute infection with cytoplasmic RNA viruses. Viral piRNA biogenesis requires ping-pong amplification by the PIWI proteins Ago3 and Piwi5. We hypothesized that Tudor proteins are required for viral piRNA production and performed a knockdown screen targeting all *Ae. aegypti* Tudor genes. Knockdown of several Tudor genes resulted in reduced viral piRNA levels, with silencing of AAEL012437 having the strongest effect. This protein, which we named Veneno, associates directly with Ago3 in an sDMA-dependent manner and localizes in cytoplasmic foci reminiscent of piRNA processing granules of *Drosophila*. Veneno-interactome analyses reveal a network of co-factors including the orthologs of the *Drosophila* piRNA pathway components Vasa and Yb, which in turn interacts directly with Piwi5. We propose that Veneno assembles a multi-protein complex for ping-pong dependent piRNA production from exogenous viral RNA.

## Introduction

In animals, three distinct small RNA-mediated silencing pathways exist: the micro (mi)RNA, small interfering (si)RNA and PIWI-interacting (pi)RNA pathways^1^. In all three, a small RNA molecule provides sequence specificity to guide a member of the Argonaute protein family to target RNA. Whereas miRNAs and siRNAs associate with proteins of the AGO clade of this family, piRNAs are loaded onto PIWI clade proteins exclusively, forming piRNA induced silencing complexes (piRISCs)^2^.

The piRNA pathway is primarily known for its role in transgenerational protection of genome integrity by silencing transposable elements in the germline^3,4^. Despite ubiquitous expression of piRNAs across metazoans, our knowledge on the molecular mechanisms that govern the piRNA pathway is limited to only a small number of model organisms^5^. In the *Drosophila melanogaster* germline, single-stranded precursors are produced from genomic piRNA clusters that contain remnants of transposable elements^6^. These precursors leave the nucleus and are processed to give rise to a pool of primary piRNAs. The PIWI proteins Piwi and Aubergine (Aub) are preferentially loaded with such primary piRNAs that bear a uridine at the first nucleotide position (1U) and are generally antisense towards transposon mRNAs^6–8^. Upon loading with a piRNA, Piwi migrates to the nucleus to enforce transcriptional silencing, while Aub targets and cleaves cognate transposon RNA in an electron-dense perinuclear structure termed *nuage*^3,9^. The 3′-fragments that remain after Aub-cleavage are subsequently loaded onto the PIWI protein Ago3 and processed further into mature secondary piRNAs, which are primarily of sense orientation^6,7^. In turn, the resulting Ago3-piRISCs can target and cleave precursor transcripts to produce new antisense Aub-associated piRNAs, thus completing the so-called ping-pong amplification cycle. As Aub preferentially binds 1U piRNAs and cleaves target RNAs between nucleotides 10 and 11, Ago3-associated secondary piRNAs mostly have adenosine residues at their tenth nucleotide position (10A). The resulting 10 nt overlap of 5’ ends and 1U/10A nucleotide biases are hallmarks of piRNA production by the ping-pong amplification loop and are referred to as the ping-pong signature^6,7^. In addition to ping-pong amplification of piRNAs, Aub- and Ago3-mediated cleavage can induce phased production of downstream Piwi-associated piRNAs which have a strong 1U preference^10,11^.

Ping-pong amplification of piRNAs was previously thought to be restricted to germline tissues, but recently, ping-pong dependent piRNA production has been demonstrated in somatic tissues of several arthropods, among which hematophagous mosquitoes of the *Aedes* family^12,13^. These anthropophilic vector mosquitoes, primarily *Ae. aegypti* and *Ae. albopictus,* are crucial for the transmission of several arthropod-borne (arbo)viruses that cause debilitating diseases such as dengue, chikungunya and Zika^14^. Since arboviral infectivity is greatly affected by the ability of the virus to replicate in the vector, mosquito antiviral immunity is a key determinant for virus transmission. Intriguingly, while causing severe disease in vertebrate hosts, arboviruses are able to replicate to high levels in the mosquito without apparent fitness cost to the insect^15^. An efficient immune response based on small interfering (si)RNAs is thought to contribute to this tolerance phenotype, as genetic interference with viral siRNA production causes elevated virus replication accompanied by increased mosquito mortality^16–19^.

In addition to siRNAs, arbovirus infection results in *de novo* production of virus-derived piRNAs (vpiRNAs) in aedine mosquitoes and cell lines, suggesting that two independent small RNA pathways contribute to antiviral immunity in these insects^13^. In *Ae. aegypti* cells, vpiRNAs from the alphavirus Sindbis virus (SINV) are predominantly produced in a ping-pong amplification loop involving the PIWI proteins Ago3 and Piwi5^20^. These proteins associate directly with vpiRNAs, which bear the distinct 1U/10A nucleotide signature indicative of ping-pong amplification. The further configuration of protein complexes responsible for vpiRNA biogenesis is currently unknown. Moreover, it is unclear whether vpiRNA production requires dedicated complexes that differ from those that mediate biogenesis of piRNAs from other substrates (e.g. transposons or host mRNAs).

Studies in *D. melanogaster* and other model organisms have shown that TUDOR domain-containing (Tudor) proteins serve important functions in piRNA biogenesis, including the prevention of non-specific degradation of piRNA substrates, facilitating PIWI protein interactions, and aiding in small RNA loading onto specific PIWI proteins^3,4,21,22^. TUDOR domains contain conserved motifs that are known to interact with symmetrically dimethylated arginines (sDMAs), a common post-translational modification on PIWI proteins^23–25^. Consequently, Tudor proteins may serve as adaptor molecules that facilitate the assembly of multi-molecular complexes involved in vpiRNA biogenesis in *Ae. aegypti.*

To test this hypothesis, we performed a functional knockdown screen of all predicted *Ae. aegypti* Tudor proteins, in which knockdown of the hitherto uncharacterized Tudor protein AAEL012437 shows the most prominent vpiRNA depletion. Because of this dramatic effect on vpiRNA biogenesis and the fact that its direct *D. melanogaster* ortholog (CG9684) is largely uncharacterized, we decided to focus our attention on this protein, which we named Veneno (Ven). Ven-depletion dramatically reduces piRNA production from both viral RNA strands, while ping-pong dependent piRNA production from endogenous sources (Ty3-gypsy transposons and histone H4 mRNA) is only mildly affected. Ven resides in cytoplasmic foci, reminiscent of piRNA processing granules in *Drosophila* and interacts directly with Ago3 through canonical TUDOR domain-mediated sDMA recognition. In addition, Ven associates with orthologs of *Drosophila* piRNA pathway components Vasa (AAEL004978) and Yb (AAEL001939) ^9,26–31^, which in turn binds Piwi5. We propose that this complex supports efficient ping-pong amplification of vpiRNAs by the PIWI proteins Ago3 and Piwi5.

## Materials and Methods

### Tudor gene identification and ortholog detection

To allow comprehensive identification of all *Ae. aegypti* Tudor genes, we combined HHpred homology detection with Jackhmmer iterative searches^32,33^. Subsequently, identified sequences were aligned using T-Coffee to determine orthologous relations between *Ae. aegypti* and *D. melanogaster* Tudor proteins^34^. In view of the length of the Tudor domain (~50 AA) and low levels of sequence conservation among the family members, neighbor joining was used to identify orthology relations, which were consistent with the domain organization of the proteins. See Supplementary Information for a detailed description of our approach.

### Transfection and infection of Aag2-cells

In knockdown experiments, cells were transfected with dsRNA and re-transfected 48 hours later to ensure prolonged knockdown. Where indicated, cells were infected with a recombinant Sindbis virus expressing GFP from a duplicated subgenomic promoter (SINV-GFP; produced from pTE3’2J-GFP^35,36^) at a multiplicity of infection (MOI) of 1 and harvested 48 hours post infection. For immunofluorescence (IFA) and immunoprecipitation (IP) experiments, Aag2 cells were transfected with expression plasmids encoding tagged transgenes and, where indicated, infected with SINV (produced from pTE3’2J^35^) at an MOI of 1 three hours after transfection. All samples were harvested 48 hours after transfection. For mass spectrometry (MS) experiments, expression plasmids were transfected into cells using polyethylenimine (PEI) and infected 24 hours later with SINV at an MOI of 0.1. MS-samples were harvested 72 hours post infection. For a more detailed description of cell culture conditions, generation of stable cell lines, generation of expression vectors, and virus production, see Supplementary Information.

### Small RNA northern blotting and RT-qPCR

For small RNA northern blotting, RNA was size separated on polyacrylamide gels and cross-linked to nylon membranes using 1-ethyl-3-(3-dimethylaminopropyl)carbodiimide hydrochloride^37^. Small RNAs were detected using ^32^P-labelled DNA oligonucleotides. For quantitative RT-qPCR analyses, DNaseI-treated RNA was reverse transcribed and PCR amplified in the presence of SYBR green. See Supplementary Information for a detailed description of the experimental procedures, sequences of probes used for northern blotting, and qPCR primers.

### Preparation of small RNA libraries and bioinformatic analyses

Total RNA from Aag2 cells transfected with dsRNA targeting either Veneno or Firefly Luciferase was used to generate small RNA deep sequencing libraries. For each condition, three transfections and library preparations were performed in parallel using Illumina’s Truseq technology, as described in^38^. See Supplementary Information for a description of the additional details on the analyses of deep sequencing libraries.

### Fluorescence and microscopy

Fluorescent imaging was performed on paraformaldehyde-fixed Aag2-cells that were permeabilized and counterstained using Hoechst-solution. Confocal images were taken using the Olympus FV1000 microscope. Images used for the quantification of GFP-signal granularity were taken using the Zeiss Axio Imager Z1 with ApoTome technology. See Supplementary Information for more details of the experimental approach.

### Immunoprecipitation and western blotting

GFP- and RFP-tagged transgenes were immunoprecipitated using GFP- and RFP-TRAP beads (Chromotek), respectively, according to manufacturer’s instructions. V5-tagged transgenes were purified using V5-agarose beads (Sigma). For Ago3 and Piwi5 immunoprecipitation (IP) experiments, antibodies targeting endogenous proteins were added to lysates at 1:10 dilution and incubated for 4 hours at 4°C, followed by overnight binding to Protein A/G PLUS agarose beads (Santa Cruz). Protein extracts were resolved on polyacrylamide gels, blotted to nitrocellulose membranes, and probed with the indicated antibodies. Details on generation of Ago3 and Piwi5 antibodies, experimental procedures, and antibody dilutions can be found in the Supplementary Information.

### Mass spectrometry

For mass spectrometry analysis, precipitated proteins were washed extensively and subjected to on-bead trypsin digestion as described previously^39^. Subsequently, tryptic peptides were acidified and desalted using Stagetips^40^ before elution onto a NanoLC-MS/MS. Mass spectra were recorded on a QExactive mass spectrometer (Thermo Scientific). For detailed experimental procedures and the analyses of mass spectra, see Supplementary Information.

### Density gradient fractionation

Lysate was separated on a 10-45% Sucrose gradient by ultracentrifugation. Subsequently, protein was precipitated in acetone and trichloroacetic acid and RNA was extracted using acid phenol/chloroform. For more details on the experimental procedures, see Supplementary Information.

## RESULTS

### Comprehensive identification of Tudor proteins in *Aedes aegypti*

Tudor proteins play fundamental roles in the biogenesis of piRNAs in both vertebrate and invertebrate species^21,22^. We therefore hypothesized that processing of viral RNA into piRNAs in *Ae. aegypti* also involves members of this protein family. To faithfully identify all *Ae. aegypti* Tudor genes and their corresponding fruit fly orthologs, we used a homology-based prediction approach combining HHPred and Jackhmmer algorithms^32,33^. First, we used HHPred homology detection to predict *D. melanogaster* TUDOR domain sequences, which were subsequently used as input for Jackhmmer iterative searches to identify all *D. melanogaster* and *Ae. aegypti* TUDOR domains. Ultimately, a neighbor joining tree was made based on a TUDOR domain alignment generated with T-Coffee^34^, which enabled the identification of orthologous relationships between *Ae. aegypti* and *D. melanogaster* Tudor proteins (Figure 1).

**Figure 1.**
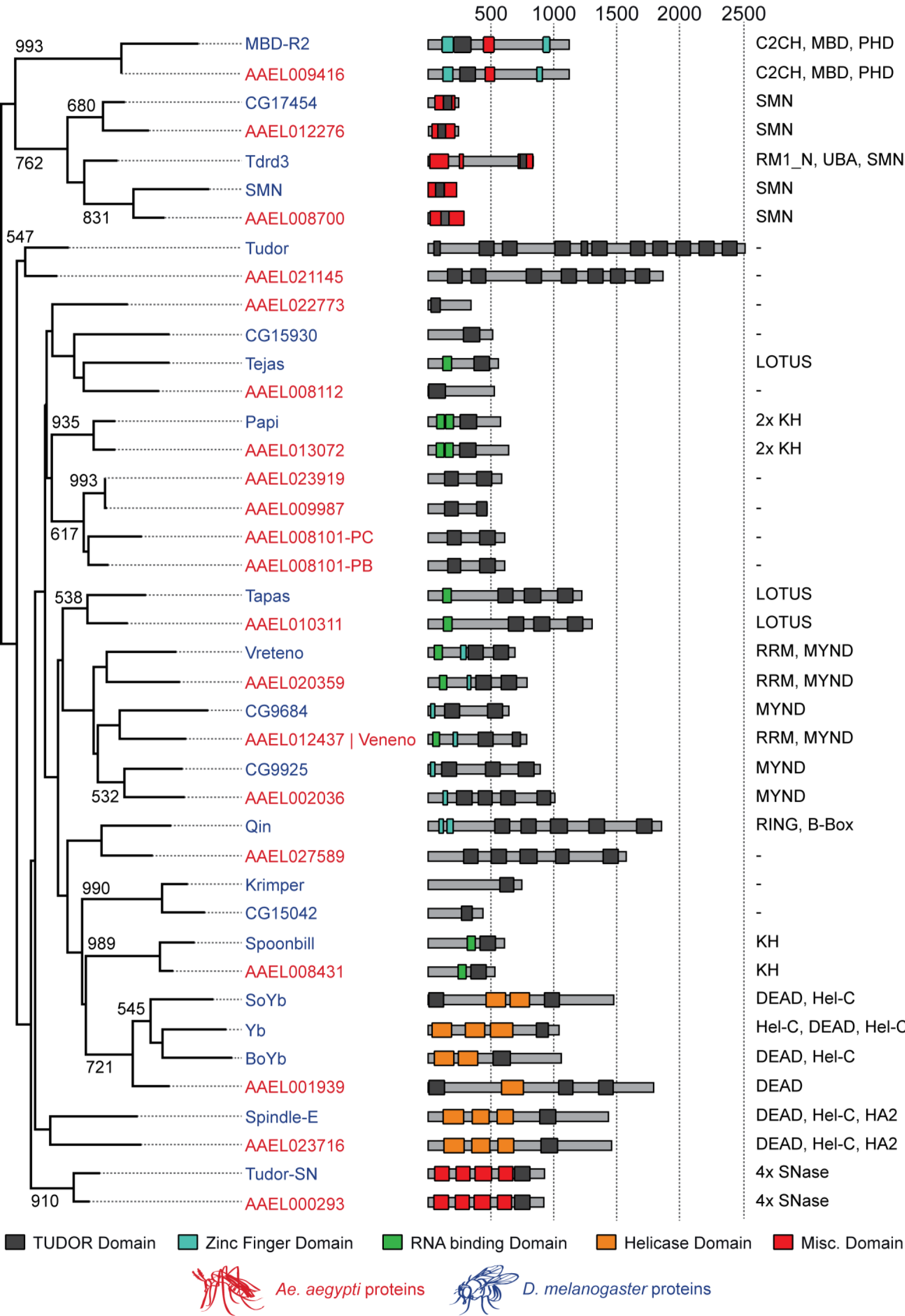
Orthologous Tudor proteins in D. melanogaster and Ae. aegypti. On the left, a neighbor joining tree based on TUDOR domains from Ae. aegypti (red) and D. melanogaster (blue) is shown. Numbers indicate bootstrap values for 1000 iterations; only values >500 are shown. In the middle, predicted domain structures of Tudor proteins are drawn schematically, with TUDOR domains shown in black, zinc fingers in blue, putative RNA binding domains in green, domains associated with helicase activity in orange, and all other domains in red. Numbers at the top indicate protein length in amino acids. On the right, protein domains other than Tudor domains are presented, ordered from amino to carboxyl terminus, as indicated in the middle panel. As the AAEL008101 gene produces two splice variants encoding Tudor domains of slightly different composition (PB and PC), we included both as separate entities in our multiple sequence alignment. B-box, B-box type zinc finger (Zf) domain; C2CH, C2CH-type Zf domain; C2H2, C2H2-type Zf domain; DEAD, DEAD box domain; HA2, Helicase-associated domain; Hel-C, helicase C domain; KH, K homology RNA-binding domain; LOTUS, OST-HTH/LOTUS domain; MBD, Methyl-CpG-binding domain; MYND, MYND (myeloid, Nervy, DEAF-1)-type Zf domain; PHD, PHD-type Zf domain; RING, RING-type Zf domain; RMI1_N, RecQ mediated genome instability domain; RRM, RNA recognition motif; SMN, survival motor neuron domain; SNase, Staphylococcal nuclease homologue domain; UBA, ubiquitin associated domain.

While the bootstrap values suggest relatively low phylogenetic signal in the TUDOR domains themselves, the majority of *Ae. aegypti* Tudor proteins cluster with a single *D. melanogaster* ortholog with a highly similar domain composition, providing independent support for the orthology relationships. Some genes however (e.g. AAEL022773, AAEL008101, AAEL009987 and AAEL023919) lack clear one-to-one orthology with *Drosophila* counterparts, suggesting that these genes emerged as a result of duplication events that occurred in the *Culicidae.* Conversely, CG15042 and Krimper in *D. melanogaster* likely resulted from a duplication in the *Drosophilidae*. Alternatively, the proteins without an ortholog may have been lost from the *Drosophila* lineage, or the proteins have diversified to an extent that they are no longer recognized as orthologous in the multiple sequence alignment. The AAEL008101 gene encodes two splice variants, of which only AAEL008101-PB is expressed in Aag2 cells (Figure S1C). Lastly, the *Ae. aegypti* genome encodes only one ortholog for the *D. melanogaster* Yb protein subfamily (Yb, SoYb and BoYb), namely AAEL001939, which we refer to as Yb. In cases where there is clear one-to-one orthology, *Aedes aegypti* will be named after their *Drosophila* ortholog throughout this study.

### *Aedes aegypti* Tudor proteins are involved in vpiRNA biogenesis

We included all identified *Ae. aegypti* Tudor proteins along with AAEL004290 in a functional knockdown screen. AAEL004290 is the ortholog of Eggless, a histone methyltransferase involved in the piRNA pathway in *D. melanogaster* which is predicted to contain TUDOR domains^41,42^, although it did not surface in our HHpred-based homology detection.

In a previous study, deep sequencing of small RNAs from Sindbis-virus (SINV) infected Aag2 cells revealed that the majority of vpiRNAs are derived from a ~200nt hotspot in the SINV-capsid gene (Figure S1A)^20^. We selected four highly abundant sense (+) strand derived vpiRNA sequences from this hotspot region for small RNA northern blotting. Knockdown of several Tudor proteins lead to reduced vpiRNA levels in Aag2 cells, with knockdown of AAEL012437 resulting in the most prominent phenotype (Figure 2). Knockdown was generally efficient, resulting in a 50 to 80% reduction of mRNA abundance for most genes (Figure 2 and S1C). For genes for which knockdown efficiency was suboptimal (Yb, AAEL008101 and MBD-R2), we performed an additional knockdown experiment using different batches of dsRNA targeting these transcripts. Here, we found that vpiRNA levels are also diminished upon knockdown of Yb and AAEL008101-RB (Figure S1B). The observed effect on vpiRNA levels cannot be explained by changes in viral replication, as only minor differences were seen in viral RNA levels across knockdowns under these experimental conditions (Figure 2 and S1B, D-E). Moreover, changes in vpiRNA production did not correlate with expression levels of capsid RNA, which is the source of vpiRNAs that we probed for in this screen (R^2^=0.0506; Figure S1F). Interestingly, histone H4 mRNA-derived piRNA production, which has previously been shown to depend on amplification by Ago3 and Piwi5^43^, was not affected by AAEL012437 knockdown (Figure 2), suggesting that this protein acts in a complex that preferentially processes piRNAs from viral transcripts. As depletion of AAEL012437 resulted in the most prominent reduction of vpiRNA levels in repeated experiments (Figure S1G-H), we proceeded with a more detailed characterization of this protein. Since the word virus comes from the Latin noun ‘poison’, we named this Tudor protein after the indicative present of the Latin verb ‘to poison’: Veneno (Ven).

**Figure 2.**
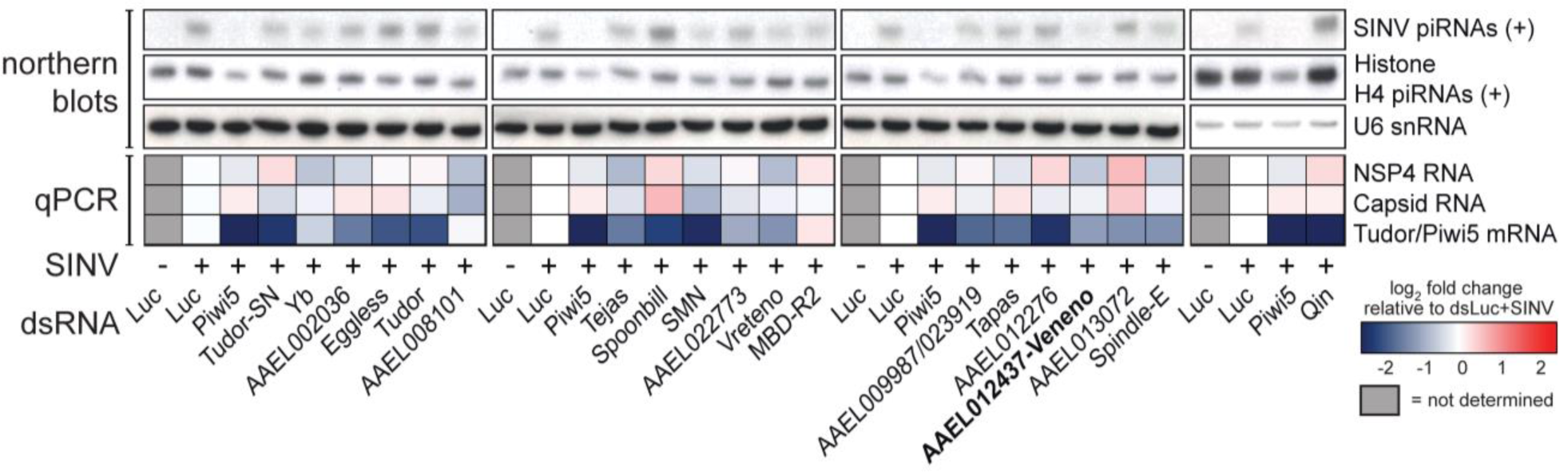
Loss of vpiRNA production upon knockdown of several Tudor proteins. Tudor genes were knocked down in Aag2 cells by dsRNA transfection after which small RNA production of (+) strand Sindbis virus (SINV) and histone H4 mRNA (H4)-derived piRNAs was assessed using northern blot analyses. As controls, dsRNA targeting luciferase (dsLuc) and Piwi5 were used as negative and positive controls, respectively. U6 snRNA was used as a loading control. Aedes proteins that have a clear ortholog with similar domain composition are named after their Drosophila orthologs. The heat map depicts relative changes in NSP4 and Capsid viral RNA abundance and Tudor/Piwi5 knockdown efficiencies as determined by RT-qPCR. All expression values were normalized to SINV-infected dsLuc control samples. Grey boxes indicate samples for which no RT-qPCR was performed.

### Depletion of Veneno predominantly affects production of viral piRNAs

Small RNA northern blotting is suitable for the detection of only a handful of highly abundant piRNAs. To enable a more comprehensive analysis of small RNA populations upon Ven knockdown (KD), we prepared small RNA deep sequencing libraries from Aag2 cells infected with SINV. Depletion of Ven resulted in a strong reduction of vpiRNA production from both strands (75% and 80% reduction of (+) and (-) strand-derived vpiRNAs, respectively), whereas viral siRNA levels were unaffected (Figure 3A-B). As reported previously^20^, SINV-derived piRNAs exhibit the 1U/10A nucleotide bias and 5’ end overlap indicative of ping-pong dependent amplification (Figure 3C-D). The vpiRNAs that remain in dsVen libraries exhibit a less pronounced nucleotide bias (Figure S2A) and 5’ end overlap (Figure 3D), indicating ping-pong amplification is reduced upon Ven-KD. The distribution of piRNAs across the viral genome and their size profile is unchanged in Ven-KD libraries (Figure S2B-E), indicating that Ven is not directly responsible for triggering vpiRNA production or determining vpiRNA length. The effect of Ven-KD on piRNA production from transposable elements is minor compared to the changes in vpiRNA production, especially for those derived from the (+) strand (25% and 55% reduction of piRNAs derived from (+) and (-) strand, respectively; Figure S3A). As the vast majority (>75%) of all transposon-derived piRNAs in our libraries originate from Ty3-gypsy elements (Figure S3B), the effects seen in this family may dominate the overall phenotype of transposon piRNAs. Stratification of transposon-derived piRNAs into subclasses indeed reveals only a mild effect of Ven-KD on piRNA production from Ty3-gypsy elements, 15% and 55% reduction of (+) and (-) strand derived piRNAs, respectively (Figure 3A). The discrepancy between the effect of Ven-KD on vpiRNA production and Ty3-gypsy-derived piRNA production is especially intriguing as Ty3-gypsy elements comprise the only major class of transposable elements that is processed into piRNAs by the ping-pong amplification loop, as is evident from their strong ping-pong signature (Figure 3C-D, S3C). Depletion of Ven results in reduced levels BEL-Pao element-derived piRNAs from both strands (Figure 3A), which make up the second-largest group of transposon-derived piRNAs (Figure S3B). In contrast to piRNAs derived from Ty3-gypsy elements, BEL-Pao piRNAs lack a 1U/10A nucleotide signature (Figure 3C) and display only a very minor 10nt overlap of 5’ends (Figure 3D). Instead, both sense and antisense BEL-Pao-derived piRNAs are enriched for 1U, suggesting that their production does not depend on ping-pong amplification but rather are on primary biogenesis or phased piRNA production. Generally, siRNA production was unchanged for all transposon subfamilies (Figure 3B, S3C). In accordance with northern blot analyses (Figure 2), ping-pong dependent histone H4 mRNA-derived piRNA levels were only mildly reduced upon Ven-KD (Figure 3A). Taken together, these findings suggest that Ven supports ping-pong dependent piRNA biogenesis preferentially from viral RNA.

**Figure 3.**
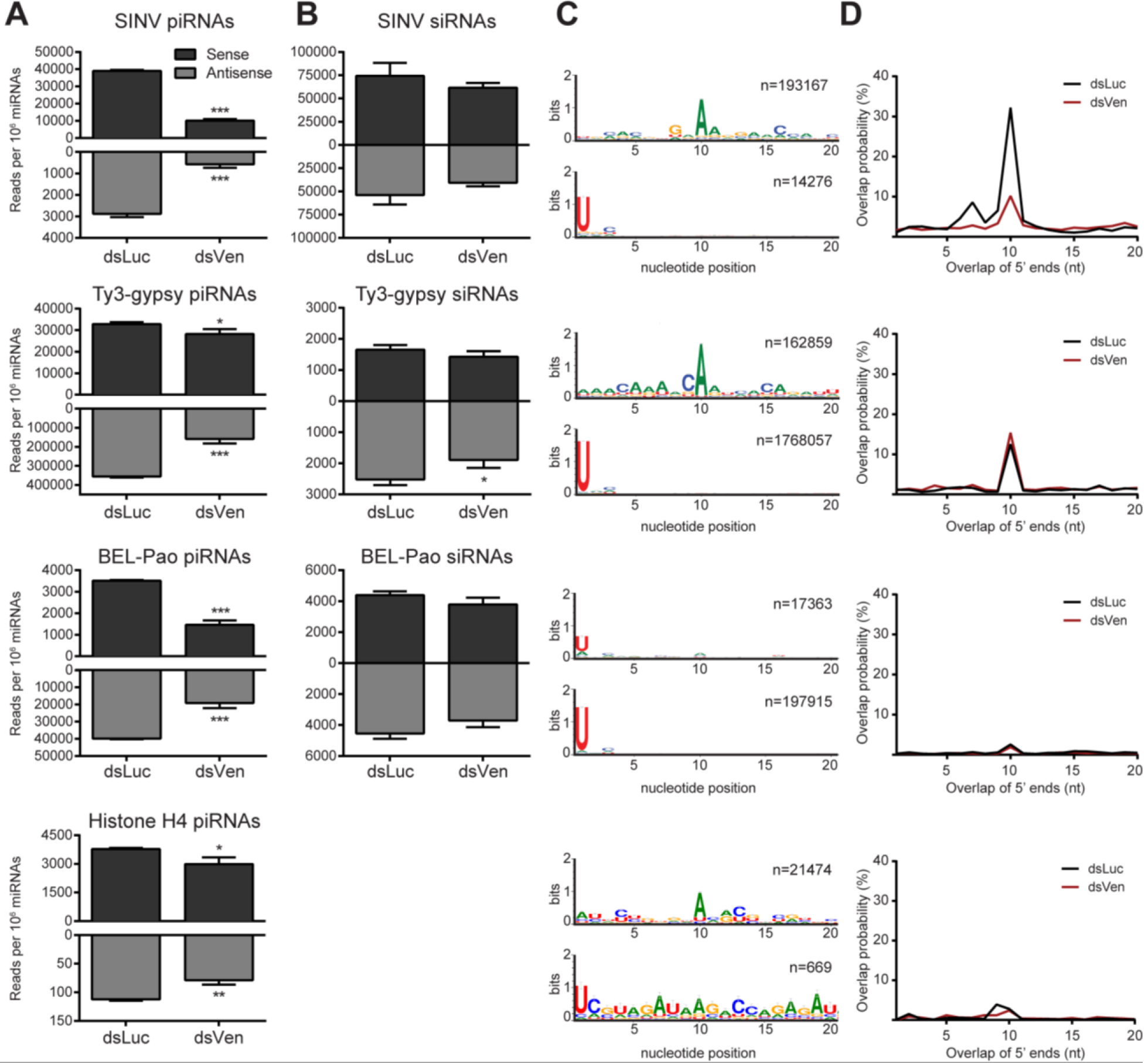
Veneno is required for efficient biogenesis of vpiRNAs. (A-B) Normalized read counts of 25-30 nt piRNAs (A) and 21 nt siRNAs (B) mapping to the Sindbis virus (SINV) genome (top row), Ty3-gypsy transposons (second row), BEL-Pao transposons (third row), and histone H4 mRNA (bottom row) upon knockdown of Veneno (dsVen) and control knockdown (Firefly Luciferase, dsLuc). Virtually no siRNA-sized reads mapping to histone H4 mRNA were found (~200 reads per library), and these are therefore not shown. (C) Nucleotide bias at the first 20 positions of the 25-30 nt small RNA reads mapping to sense strand (upper panel) and antisense strand (lower panel) of the indicated RNA substrates in dsLuc libraries (n = number of reads). (D) The probability of 5’ overlap between piRNAs from opposite strands in dsLuc and dsVen libraries for piRNAs mapping to indicated RNA substrates. For bar charts in A and B, read counts of three independent libraries were normalized to the amount of miRNAs present in those libraries and analyzed separately for the sense (black) and antisense (grey) strands. Bars indicate mean +/-standard deviation. Two-tailed student’s t-test was used to determine statistical significance (* P < 0.05; ** P < 0.01, *** P < 0.001). To generate sequence logos and 5’ overlap probability plots shown in C and D, reads of three independent libraries were combined.

### Veneno localizes to cytoplasmic foci

To further characterize the molecular function of Ven during vpiRNA biogenesis, we expressed GFP-tagged Ven and several domain mutants (Figure 4A) in Aag2 cells. In the process of cloning these constructs, we noticed that the annotation of the Ven-gene in AaegL3.5 on VectorBase was erroneous. We used Sanger sequencing of PCR products to revise the current gene annotation (Figure S4), which was corroborated by published Aag2 transcriptome data and the recently released *Aedes aegypti* mosquito reference genome assembly AaegL5^44,45^. GFP-tagged Ven accumulated in cytoplasmic foci reminiscent of the piRNA processing granules *nuage* and Yb bodies in *D. melanogaster* (Figure 4B)^9,29^. We tentatively term these foci Ven-bodies. In our revised annotation, Ven contains an RNA recognition motif (RRM) at its amino terminus. A mutant in which this motif has been removed (C91) retains its localization in Ven-bodies (Figure 4C), suggesting that putative RNA-binding by this domain is not required for granule formation. Additionally, Ven contains a Zn-finger of the MYND-type, a class of Zn-fingers predominantly involved in protein-protein interaction^46^. Removal of this MYND-domain (C234) abolishes granular accumulation of Ven (Figure 4E-F). Intrinsically disordered sequences have recently been shown to mediate RNA binding and regulate RNA metabolism^47^. Ven contains such an asparagine (N)-rich region directly upstream of the MYND-domain. A mutant in which this N-rich stretch as well as the RRM are removed (C199) but the MYND-domain is maintained, retains its localization in Ven-bodies (Figure 4D), which lends further support to the importance of the MYND-domain for the granular localization pattern.

**Figure 4.**
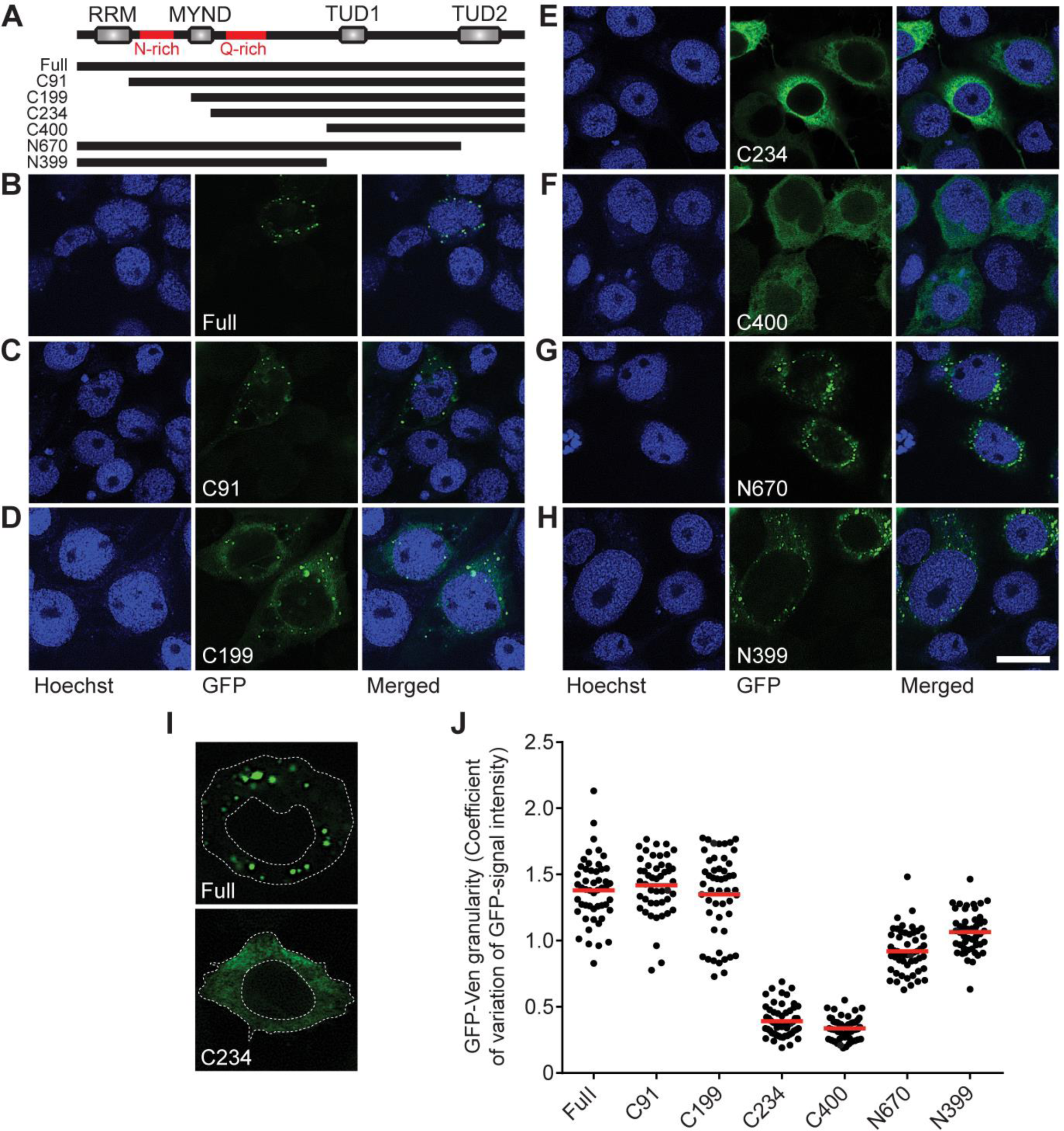
Veneno accumulates in cytoplasmic Ven-bodies. (A) Schematic representation of Ven transgenes used in immunofluorescence experiments. (Full: amino acid [aa] 1-785; C91: aa 91-785; C199: aa 199-785; C234: aa 234-785; C400: aa 400-785; N670: aa 1-670; N399: aa 1-399; red lines indicate sequences of low amino acid complexity, rich in asparagines [N] or glutamines [Q]). (B-H) Representative confocal images of Aag2 cells expressing transgenes drawn schematically in (A). Scale bar represents 10μm. (I) The cytoplasm of 46-56 individual cells expressing GFP-tagged transgenes was traced as depicted and the mean and standard deviation of signal intensity was determined to calculate the coefficient of variation as a measure of signal granularity. (J) Scatter dot plot shows the GFP-signal granularity for individual cells; the red line indicates the mean.

Similarly, a Q-rich sequence directly downstream of the MYND-type Zn-finger is not sufficient for Ven-accumulation, as removal of the MYND domain alone (C234) disrupts Ven-body formation. Upon removal of the C-terminal (N670) or both TUDOR domains (N399), Vens distinct subcellular localization is largely retained, suggesting that TUDOR domains do not play a major role in granule formation (Figure 4G-H). To allow a more comprehensive analysis of Ven-body localization across mutants, we traced the cytoplasmic GFP-signal of ~50 cells per transgene (as shown in Figure 4I) and quantified the coefficient of variance of this signal as a measure for granularity. This analysis confirms that Ven-body accumulation is abolished upon removal of the MYND-type Zn-finger. Also, a slight decrease in granularity is seen upon removal of either one (N670) or both (N399) Tudor domains (Figure 4J), suggesting that additional protein-protein interactions may stabilize the Ven-body. Altogether, these findings suggest the MYND-type Zn-finger enables Ven localization into specific Ven-bodies where additional components of the mosquito piRNA biogenesis machinery may be recruited for efficient piRNA production.

### Veneno provides a molecular scaffold for a ping-pong amplification complex

As Ven is important for efficient production of ping-pong dependent vpiRNAs (Figures 2–3), we hypothesized that the protein may serve as a molecular scaffold to facilitate an interaction between the ping-pong partners Ago3 and Piwi5. To investigate this hypothesis, we immunoprecipitated GFP-tagged Ven and probed using antibodies recognizing endogenous Ago3 and Piwi5 (Figure S5A for antibody characterization). We found that Ven interacts with Ago3, but not Piwi5, regardless of an ongoing SINV-infection (Figure 5A). Immunoprecipitation of GFP alone does not copurify Ago3, confirming that the interaction is indeed mediated by Ven. To further dissect the multimolecular network in which Ven participates, we employed quantitative mass spectrometry of immunoprecipitated GFP-Ven complexes from both uninfected and SINV-infected Aag2 cells. These data confirm the association with Ago3 and reveal interesting additional Ven-interactors (Figure 5B-C and Supplementary Table 1). Specifically, Yb is enriched in Ven-complexes immunoprecipitated from both mock- and SINV-infected cells. Probing the Ven-interactome for orthologs of factors involved in the ping-pong amplification loop in *Drosophila,* we found a slight enrichment of AAEL004978, the *Ae. aegypti* ortholog of Vasa (Figure 5B-C). *Drosophila* Vasa recruits PIWI proteins to accommodate ping-pong amplification and is believed to be expressed exclusively in germline tissues. Yet, we verified that *Ae. aegypti* Vasa and other components of the piRNA biogenesis machinery are expressed in both the germline and somatic tissues in female *Ae. aegypti* mosquitoes (Figure S5B), implying the complex is capable of producing vpiRNAs upon arbovirus infection in the soma. We verified abovementioned interactions by co-purifying the constituents of the complex in reciprocal IPs followed by western blot (Figure 5D). Interestingly, we also detect Piwi5 as a direct interaction partner of Yb (Figure 5D). In sucrose density gradient fractionation, Ven co-sediments with Ago3 and Piwi5 (most prominently in fractions 4-5) and with piRNAs produced from the S-segment of the Phasi Charoen-like bunyavirus (Figure 5E), a known contaminant of the Aag2 cell line which has previously been reported to produce piRNAs through ping-pong amplification^48,49^. Altogether, this further suggests that Ven forms a multiprotein complex with Ago3 and Piwi5. Additionally, we find that Ven, Ago3, Piwi5, Yb and Vasa colocalize to Ven bodies, suggesting these granules are indeed the sites of vpiRNA biogenesis (Figure S5D). The effect of Yb-KD mirrors the effects seen in Ven-KD libraries, with the strongest reduction in vpiRNA levels and only moderate effects on transposon- and histone H4 mRNA- derived piRNAs (Figure S5E). The effects seen upon Yb-KD are less pronounced than those in Ven-KD libraries, which is likely due to relatively inefficient knockdown of Yb. Together, these findings reveal the presence of a multi-molecular complex in which the ping-pong partners Ago3 and Piwi5 are brought together by the Tudor proteins Ven and Yb to promote efficient piRNA production (Figure 6E).

**Figure 5.**
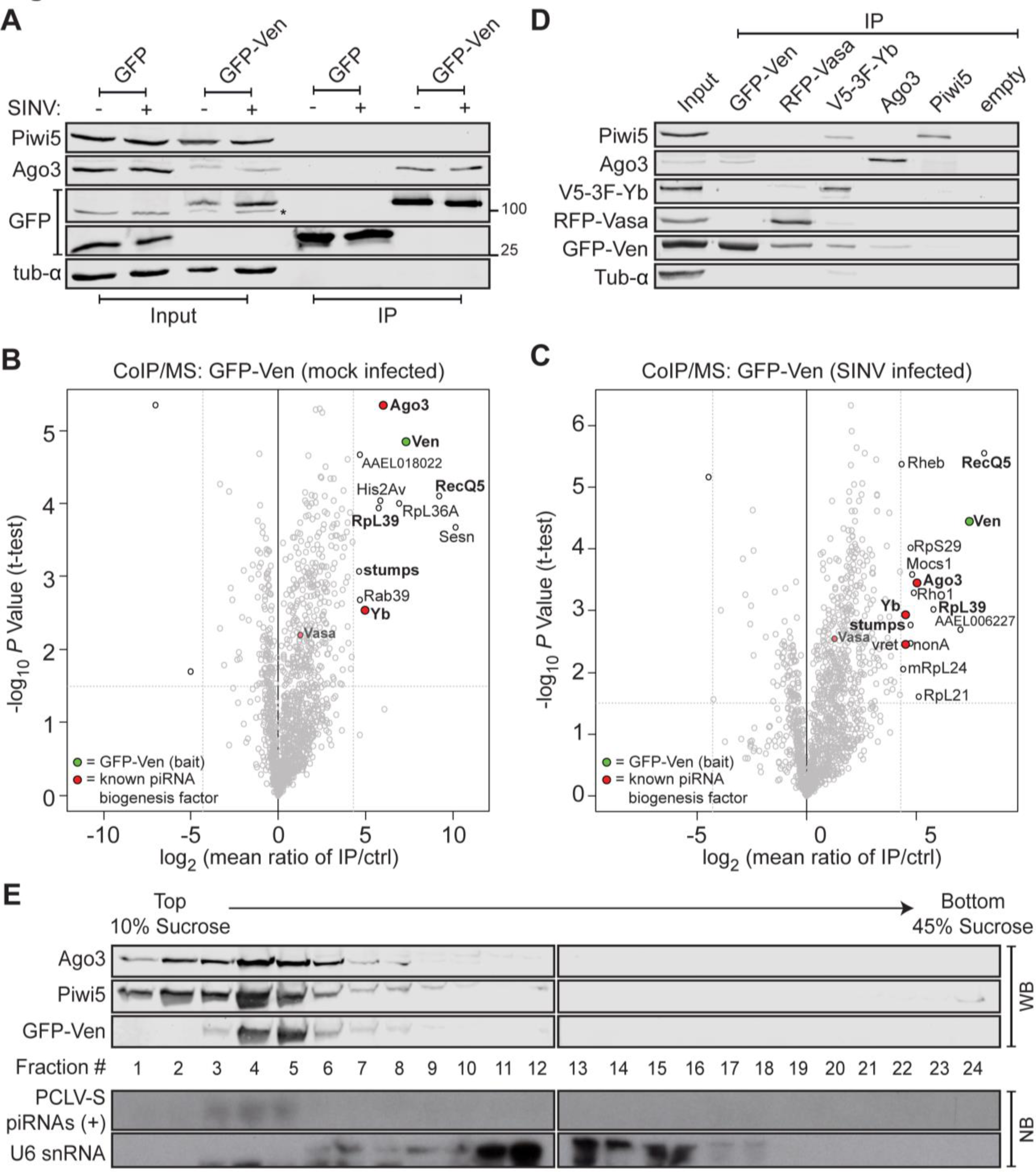
Characterization of a multi-protein complex containing the ping-pong partners Ago3 and Piwi5. **(A)** Protein lysates from SINV-infected (%) and uninfected (-) Aag2 cells transfected with expression plasmids encoding GFP or GFP-Ven before (Input) and after GFP-immunoprecipitation (IP) were analyzed for (co)purification of endogenous Ago3 and Piwi5, as well as the GFP-tagged transgene by western blot. The asterisk indicates a non-specific band. **(B-C)** Identification of Ven-interacting proteins in lysates from both mock-(B) and Sindbis virus (SINV)-infected Aag2 cells (C) by label-free quantitative (LFQ) mass spectrometry. Permutation-based FDR-corrected t-tests were used to determine proteins that are statistically enriched in the Ven-IP. The LFQ-intensity of GFP-Ven IP over a control IP using the same lysate and non-specific beads (log_2_-transformed) is plotted against the -log_10_ *P* value. Interactors with an enrichment of log_2_ fold change > 4.3; -log_10_ *P* value > 1.5 are indicated. Proteins in the top right corner represent the bait protein in green (Ven) and its interactors. Orthologs of known piRNA biogenesis factors in *D. melanogaster* are indicated in red and interacting proteins present in both mock- and SINV-infected pulldowns are shown in **bold** font. Where available, interacting proteins were named according to their ortholog in *D. melanogaster.* In case of uncharacterized orthologous *Drosophila* proteins, we assigned the Vectorbase GeneID to the protein. (**D**) Reciprocal IPs of GFP-Ven, RFP-Vasa, V5-3xflag-Yb, Ago3 and Piwi5 using antibodies targeting GFP, RFP, V5, Ago3 and Piwi5, respectively. Samples were probed with antibodies against GFP, RFP, Flag, Ago3, Piwi5 and α-tubulin, as indicated. (**E**) Lysate from Aag2 cells stably expressing GFP-Ven was fractionated on a 10-45% Sucrose gradient. Protein fractions were size separated and stained using antibodies against GFP, Ago3 and Piwi5. RNA samples from those fractions were analyzed by northern blot analysis, using probes targeting abundant (+) strand ping-pong dependent piRNAs produced from the S-segment of the PCLV bunyavirus and U6 snRNA. All fractions contain proteinaceous material as is evidenced by silver staining (Figure S5D); spliceosomal ribonucleoprotein complexes are enriched in fractions 11-16.

**Figure 6.**
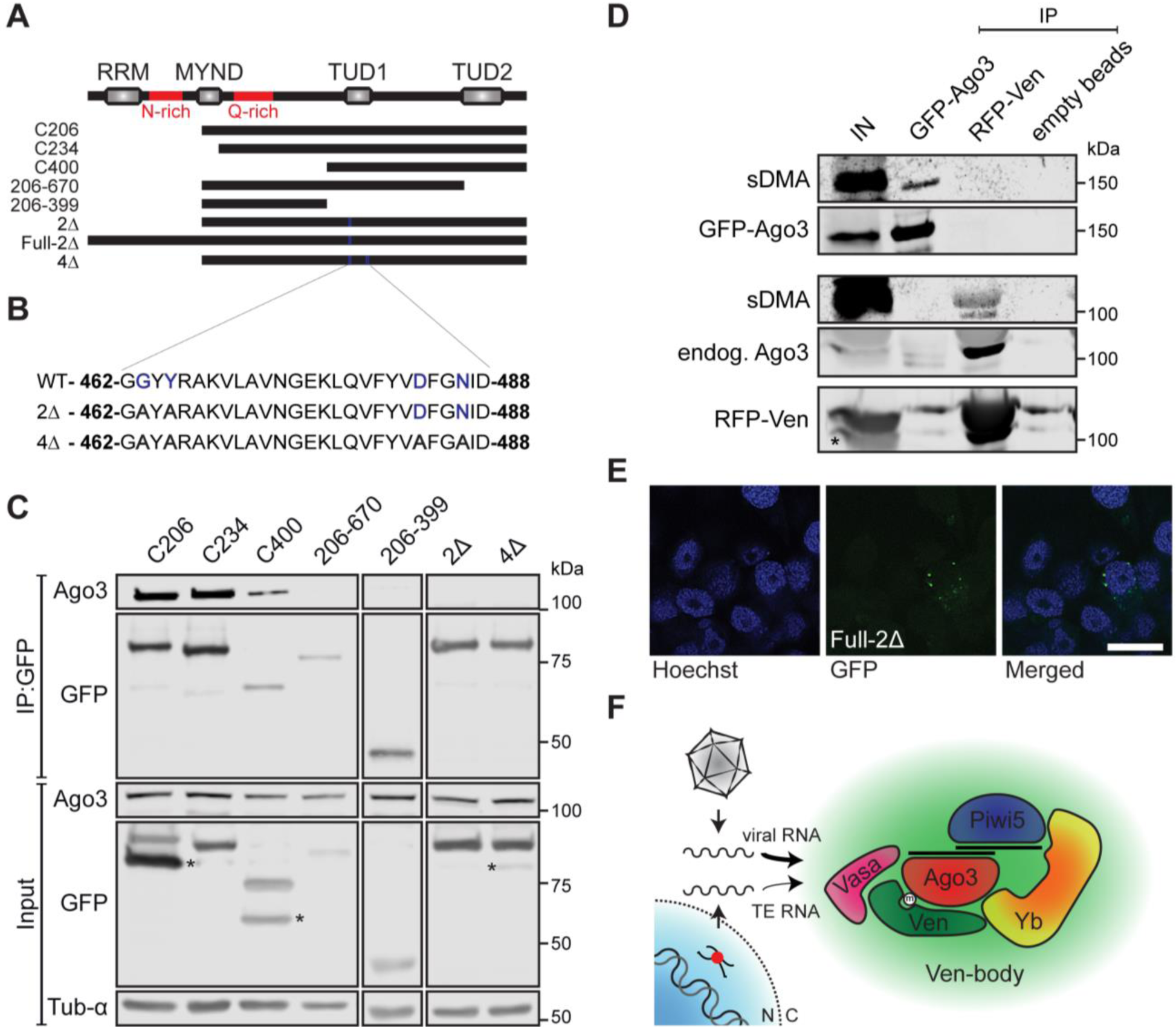
Ven-Ago3 interaction is mediated by sDMA-recognition. (**A**) Schematic representation of Veneno transgenes used in Ago3 co-IP experiments. (C206: amino acid [aa] 206-785; C234: aa 234-785; C400: aa 400-785; 206-669: aa 206-669; 206-399: aa 206-399; 2Δ: C206-G463A/Y465A; Full-2Δ: G463A/Y465A; 4Δ: C206-G463A/Y465A/D483A/N486A; red lines indicate sequences of low amino acid complexity, rich in asparagines [N] or glutamines [Q]). (**B**) Sequence corresponding to a part of the first Tudor domain with residues indicated that were mutated in the 2Δ and 4Δ transgenes. Residues indicated in blue are predicted to be involved in sDMA recognition. (**C**) Lysates from Aag2 cells expressing indicated GFP-tagged Ven transgenes were subjected to GFP-IP and subsequently analyzed for co-purification of Ago3 by western blot. α-Tubulin serves as loading control. Asterisks indicate non-specific bands. (**D**) Lysate from Aag2 cells stably expressing GFP-Ago3 and transiently transfected with a plasmid encoding RFP-Ven was immunoprecipitated using GFP-, RFP- and empty beads. Western blots were stained using antibodies for GFP, RFP, and symmetrical dimethylated arginines (sDMA). The asterisk indicates a non-specific band. (**E**) Representative confocal image of Aag2 cells expressing GFP-tagged Ven-2Δ-mutant; scale bar represents 10μm. (**F**) Schematic model of the identified multi-protein complex responsible for ping-pong amplification of exogenous (viral) and endogenous (transposable element, TE)-derived piRNAs. The thickness of the arrows reflects the relative contribution of the complex to processing of different RNA substrates. N, nucleus; C, cytoplasm.

### Ven-Ago3 interaction depends on sDMA-recognition

Interaction between TUDOR and PIWI proteins generally depends on recognition of symmetrically dimethylated arginines (sDMAs) on PIWI proteins by an aromatic cage encoded in the TUDOR domain^25^. To further characterize the domain required for the interaction between Ven and Ago3, we immunoprecipitated truncated Ven transgenes and assessed copurification of Ago3. A truncated Ven-mutant lacking the RRM (C206-Figure 6A) still strongly associates with Ago3 (Figure 6C). Moreover, this mutant retains its association in a complex involving Yb, Vasa and Piwi5 as shown by mass spectrometry (Figure S6A, Supplementary Table 2) and reciprocal IPs (Figure S6B). The MYND-domain mutant (C234), in which the distinct localization pattern is distorted (Figure 4), still aptly binds Ago3, suggesting that granular localization in Ven-bodies is not required for Ago3-interaction (Figure 6C). The carboxyl terminus containing two TUDOR domains (C400) is sufficient for interaction with Ago3, whereas Ago3-binding is lost upon deletion of the second (206-669) or both (206-399) Tudor domains (Figure 6C). However, this loss of binding may result from reduced expression or stability of these mutants. Hence, to further specify whether Ven-Ago3 interaction is TUDOR domain mediated, we generated Ven transgenes carrying point mutations in residues predicted to be involved in sDMA recognition (2Δ: C206-G463A/Y465A and 4Δ: C206-G463A/Y465A/D483A/N486A; Figure 6B). We found that the second TUDOR domain of Ven is atypical in that only one of the predicted aromatic cage residues is conserved (Figure S6C). We therefore analyzed binding of Ago3 to Ven that carries point mutations in the first TUDOR domain. Interaction with Ago3 was lost in these mutants, suggesting that the first TUDOR domain of Ven binds Ago3 in a canonical sDMA-dependent manner (Figure 6C). It is likely that the C-terminal TUDOR domain is not involved in Ago3 binding via sDMAs since critical residues are not conserved (Figure S6C). However, we cannot fully exclude that cooperative binding of both TUDOR domains by Ago3 is required for efficient association with Ven. To verify that Ago3 bears sDMA modifications, we made use of Aag2 cells stably expressing GFP-tagged Ago3 to enable simultaneous detection of GFP-Ago3 and sDMA modifications. We found a specific sDMA signal overlapping with the signal of immunopurified GFP-Ago3 (Figure 6D), indicating Ago3 indeed contains symmetrically dimethylated arginines. Endogenous Ago3 present in Ven-complexes also bears symmetrically dimethylated arginines, further supporting the notion that Ven-Ago3 interaction is mediated by sDMA recognition. The interaction with Ago3 is not required for localization of Ven in Ven-bodies, as introducing the indicated point mutations (G463A/Y465A) in the context of the full length protein does not affect its subcellular localization pattern (Figure 6E). Altogether, our findings support a model in which Veneno, through sDMA recognition, recruits Ago3 to a multi-molecular complex that promotes ping-pong amplification of piRNAs preferentially from exogenous RNAs (Figure 6F).

## DISCUSSION

Mosquito antiviral immunity largely relies on the processing of viral dsRNA into virus-derived siRNAs that direct the degradation of viral RNA. The discovery of *de novo* production of vpiRNAs from arboviral RNA however, uncovered the intriguing possibility of an additional small RNA-based line of defense against arboviruses. Processing of viral dsRNA into vsiRNAs by the siRNA pathway has been thoroughly characterized in mosquitoes^50,51^. As of yet, it is unclear how viral RNA produced in the cytoplasm is entered into the piRNA pathway, especially as canonical substrates for the piRNA pathway are genomically encoded single-stranded precursors^3,4^. To better understand how viral RNA is detected by the mosquito piRNA pathway, more insights into the multimolecular machinery that processes viral RNA into vpiRNAs is needed.

In *Ae. aegypti,* vpiRNAs are amplified by the ping-pong partners Ago3 and Piwi5, but auxiliary proteins involved in this process were unknown. As a tightly regulated network of Tudor proteins promotes production of piRNAs in *Drosophila*^21,22^, we performed a comprehensive knockdown screen to evaluate the role of *Ae. aegypti* Tudor proteins in vpiRNA biogenesis. Knockdown of several Tudor genes affects vpiRNA biogenesis, with knockdown of Veneno (Ven) resulting in the strongest depletion of vpiRNAs. Additional candidates that show depletion in vpiRNA levels are Yb and AAEL008101; a gene which does not contain a one-to-one ortholog in the fruit fly. Whereas involvement of Yb in vpiRNA biogenesis is likely explained by its central place in the multi-protein complex discovered in this study, the molecular function of AAEL008101 remains to be elucidated. We cannot exclude that additional Tudor proteins play a role in vpiRNA biogenesis, which may be masked by redundancy of paralogous proteins or residual protein activity after suboptimal knockdown efficiency.

Thus far, the direct ortholog of Ven in *D. melanogaster* (CG9684) has not been studied extensively. In a systematic analysis of all *Drosophila* Tudor proteins, germline-specific knockdown of CG9684 did not affect steady-state levels of transposon transcripts or female fertility rate^28^. This study, however, did not evaluate the effect of CG9684 knockdown on small RNA populations.

Ven accumulates in cytoplasmic foci similar to piRNA processing bodies in the fly. In *Drosophila* somatic follicle cells, which surround the germ cells, primary piRNA biogenesis takes place in Yb bodies. One of the core factors present in these structures is their eponym Yb^28–30^. Yet, no piRNA amplification takes place in Yb bodies, since the ping-pong partners Aub and Ago3 are not expressed in follicle cells^52,53^. In contrast, in *Drosophila* germ cells piRNA amplification takes place in the *nuage* and one of the core proteins of this perinuclear structure is the helicase Vasa^9,26,27^. In *Drosophila* and silkworm, Vasa is directly implicated in secondary piRNA amplification by preventing non-specific degradation of piRNA precursors and facilitating their transfer to PIWI proteins^26^. Yb is not present in *nuage* but it has been suggested that its function may be taken over by its paralogous family members: brother and sister of Yb (BoYb and SoYb, respectively)^28^. In *Ae. aegypti* only one paralog of Yb is encoded, which associates directly with Ven and Piwi5. The presence of a multi-protein complex containing orthologs of Vasa and Yb supports the idea that Ven-bodies resemble *nuage-like* piRNA processing bodies. Similar to Ven, the *Drosophila* Tudor protein Krimper localizes in perinuclear granules, which are lost upon deletion of the amino terminus of the protein^54,55^. While Krimper directly interacts with both partners in the ping-pong loop in flies (Ago3 and Aub), Ven associates exclusively with Ago3. Moreover, while Krimper-Ago3 interaction is retained when using an arginine-methylation-deficient mutant of Ago3 in fruit flies, an sDMA-recognition-deficient mutant of Ven is unable to bind Ago3 in *Ae. aegypti.* Thus, sDMA modifications seem to be required for Ven-Ago3 interaction in mosquitoes, but dispensable for Krimper-Ago3 association in fruit flies.

Knockdown of Ven greatly affects production of piRNAs derived from exogenous viral RNA, while only a modest reduction of endogenous histone H4 mRNA- and transposon-derived secondary piRNA levels is seen. The apparent stability of histone H4 derived piRNAs is especially surprising, as their production has previously been shown to depend on ping-pong amplification involving the PIWI proteins Ago3 and Piwi5^43^. Similarly, ping-pong dependent piRNA production from Ty3-gypsy transposable elements is affected only mildly by Ven-KD.

Bel-Pao-derived piRNA production is largely independent of ping-pong amplification, as is evident from the weak 1U/10A signature and 10nt overlap of piRNA 5’ends. Therefore, we were surprised to find a strong reduction in Bel-Pao-derived piRNAs upon Ven-KD. Apart from secondary piRNA production in the ping-pong loop, piRNA-mediated cleavage of transposon mRNA may trigger the production of phased piRNAs bearing a strong 1U bias^10,11^. This mechanism of phased piRNA production seems to be particularly active in *Ae. aegypti*^56^. Hence, a modest reduction of ping-pong dependent piRNA levels may result in strong reduction of phased piRNA production which could explain the strong effect of Ven-KD of production of BEL-Pao-derived piRNAs.

Our data suggest that Ven is involved in specifying the substrate for piRNA production and may preferentially shuttle viral RNA into the ping-pong loop. It would be interesting to assess whether viral RNA from other arbovirus families are similarly affected by Ven knockdown, which would point towards a more general role of Ven in self-nonself discrimination. Dependency on specific co-factors for the biogenesis of small RNAs from different RNA sources is not unprecedented. For example, the siRNA pathway co-factor Loqs-PD is required for processing of endogenous siRNA-precursors, but is dispensable for siRNA production from exogenous dsRNA or viral RNA^57,58^. Another study showed that the Tudor protein Qin/Kumo specifically prevents (+) strand transposon RNAs from becoming Piwi-bound piRNAs during the process of piRNA phasing^59^. Analogies can also be drawn to the vertebrate piRNA pathway, where Tdrd1, the closest mouse ortholog of Ven, ensures processing of the correct transcripts by the piRNA pathway^60^. Accordingly, the PIWI protein Mili contains a disproportionally large population of piRNAs derived from cellular mRNA and ribosomal RNA in *Tdrd1* knockout mice. In a similar fashion, Ven could promote processing specifically of viral RNA by the mosquito piRNA pathway. Yet, we expect the molecular mechanism underlying this Tudor protein-guided sorting to be different as Tdrd1 interacts with Mili, the PIWI protein that predominantly binds 1U biased primary piRNAs, whereas Ven associates with Ago3, which mainly binds 10A biased secondary piRNAs.

A sophisticated network of accessory proteins that guides diverse RNA substrates into distinct piRISC complexes may be of particular importance in *Ae. aegypti* as this mosquito species encodes an expanded PIWI gene family consisting of seven members^61,62^, of which four (*Ago3* and *Piwi 4-6*) are expressed in somatic tissues^63^. Moreover, the repertoire of RNA molecules that are processed into piRNAs is extended to include viral RNA^13^. Tudor proteins like Veneno may therefore aid in streamlining piRNA processing and allow flexible adaptation of the piRNA pathway in response to internal and external stimuli such as arbovirus infection.

## Acknowledgements

We thank members of the Van Rij laboratory for fruitful discussions, Eugene Berezikov (European Research Institute for the Biology of Aging) for bioinformatic advice, the Microscopic Imaging Centre of Radboudumc for support for confocal microscopy, and Siebe van Genesen for assistance with density gradient fractionation. Sequencing was performed by the GenomEast platform, a member of the “France Génomique” consortium (ANR-10-INBS-0009). We would also like to thank Gorben Pijlman (University of Wageningen) for kindly providing the pPUbB-GW vector, Frank van Kuppeveld (Utrecht University) for the rabbit-anti-GFP antibody, and the Carnegie Institution of Washington for the Drosophila Gateway Vector collection. This work is financially supported by a PhD fellowship from Radboud University Medical Center to PM, a European Research Council Consolidator Grant under the European Union’s Seventh Framework Programme (grant number ERC CoG 615680) and a VICI grant from the Netherlands Organization for scientific Research (grant number 016.VICI.170.090) to RPvR.

